## Supplementary material for "An auxin-inducible, GAL4-compatible, gene expression system for *Drosophila*": McClure et al - Supplementary material

[2] Present address: Queen's University Belfast, School of Biological Sciences, Belfast, BT9 5AH, UK

[3] Present address: Department of Clinical and Experimental Epilepsy, UCL Queen Square Institute of Neurology, London, United Kingdom.

### Supplementary information

#### Contents:

**Figure 1 – figure supplement 1**. Plasmid map of the AGES plasmid

**Figure 1 – figure supplement 2**. Sequence of the AGES plasmid – Separate file:

*pattB-tubP-AtTIR1-P2A-miniAID-Gal80-miniAID-SV40 - sequence.gb*

**Figure 2 – figure supplement 1**. AGES effectively induces GAL4 activity in *Drosophila* adult males

**Figure 2 - source data 1.xlsx** – separate file

**Figure 3 - source data 1.xlsx** – separate file

**Figure 4 – figure supplement 1.** AGES allows induction of pan-neuronal GAL4 activity in the adult brain

**Figure 5 - source data 1.xlsx** – separate file

**Figure 6 – figure supplement 1.** Dose-dependent NAA effects on behaviour of AGES parental controls

**Figure 6 – figure supplement 2.** Effects of AGES and GeneSwitch induced expression of Kir2.1 in PDF+ clock neurons on circadian period length and average 24h locomotor activity.

**Figure 6 - source data 1.xlsx** – separate file

**Fly food recipe**

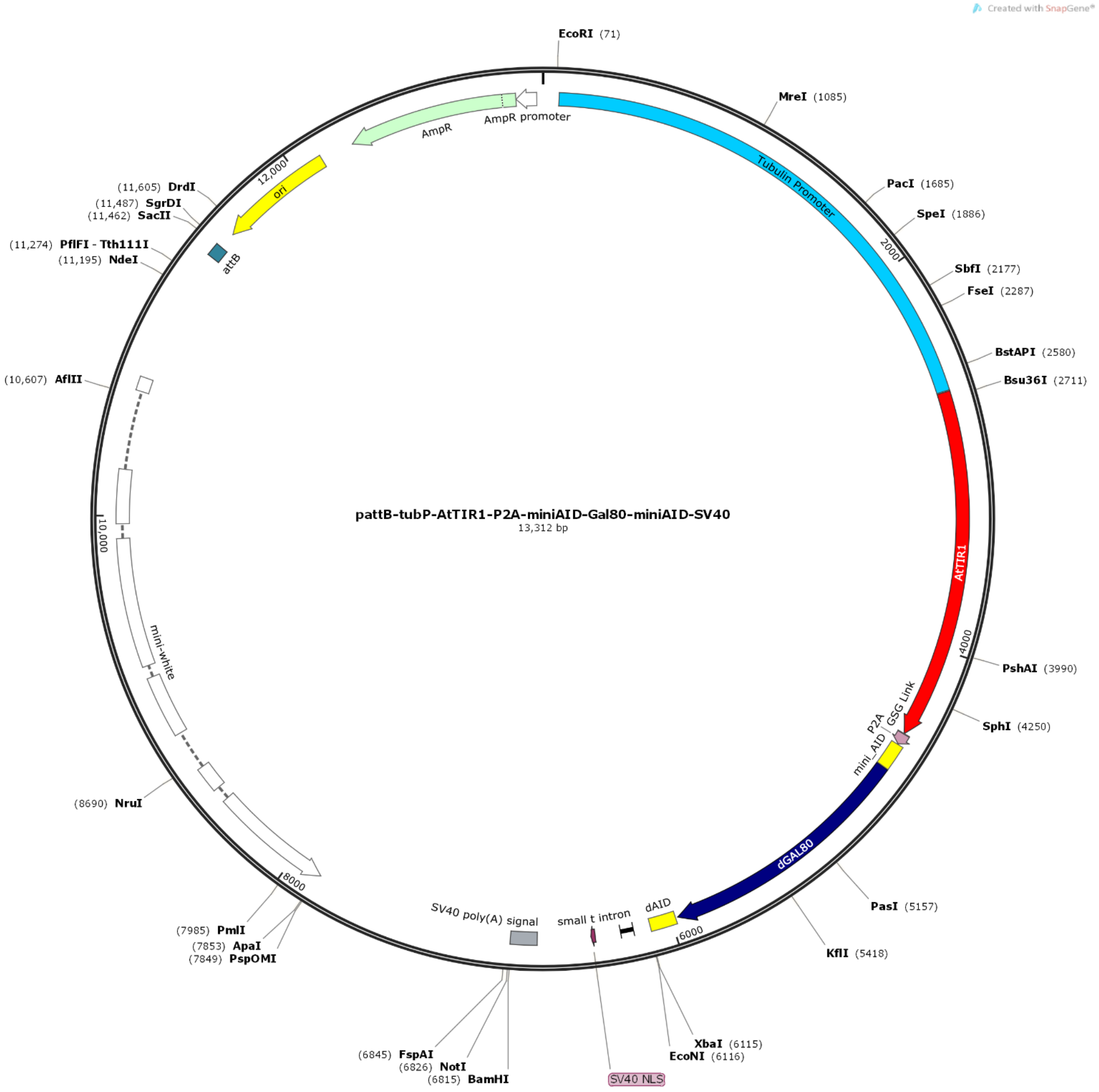


**Figure 1 – figure supplement 1. Plasmid map of the AGES plasmid**. Map of the *pattB-tubP-AtTIR1-P2A-miniAID-Gal80-miniAID-SV40* plasmid (generated using Snapgene). The full sequence is in Figure 1 – figure supplement 2.

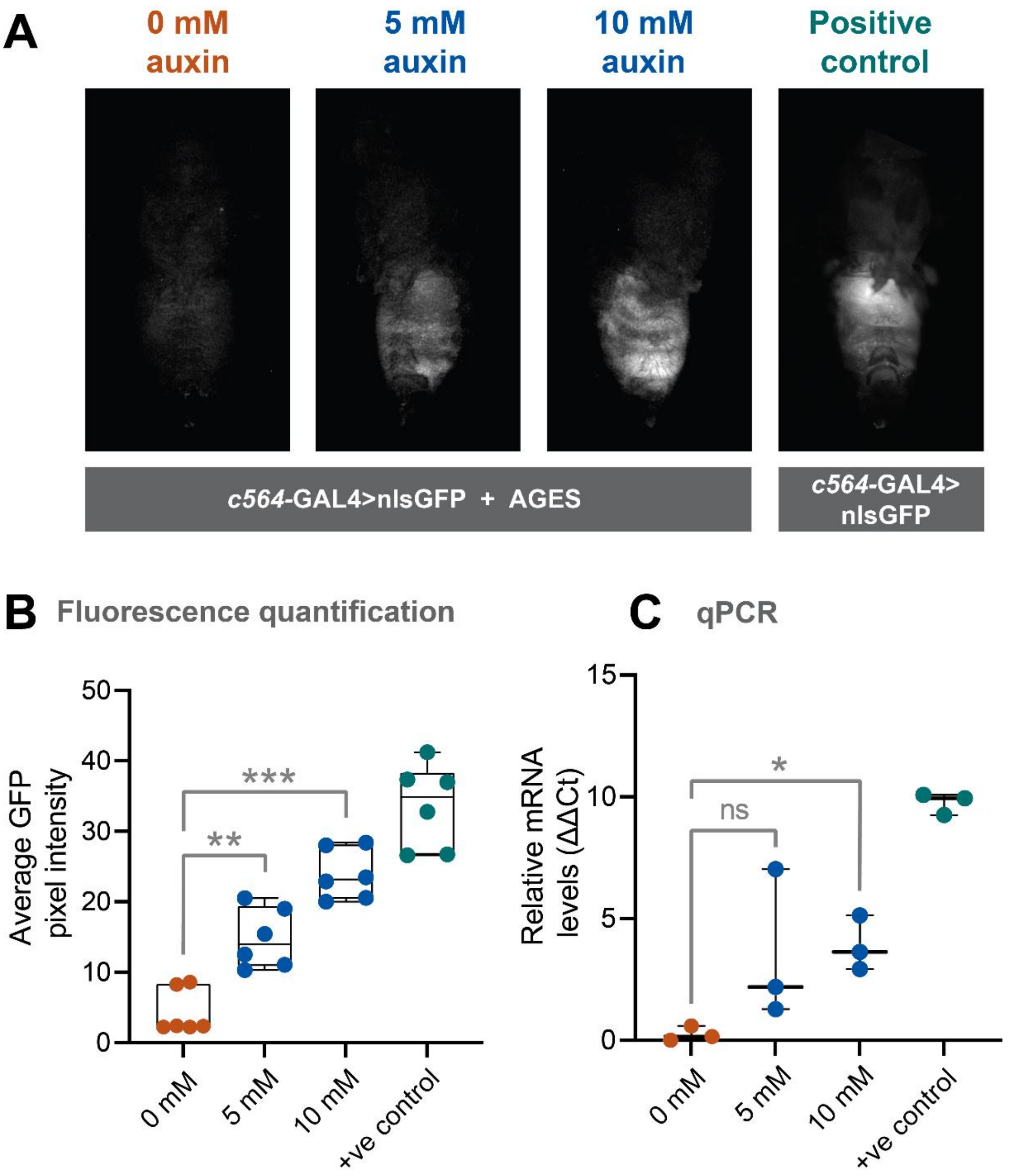


**Figure 2 – figure supplement 1. AGES effectively induces GAL4 activity in *Drosophila* adult males**. A) Ventral images of live males that express GAL4 in fat body tissue. Ingestion of food containing auxin (24 hours) induces GAL4 activity and the expression of GFP. B) Quantification of GFP levels (from 6 male abdomens). Pixel intensity thresholding was performed to isolate abdomens as regions of interest. The average pixel intensity intensities were quantified and analysed using Ordinary one-way ANOVA (**, $p = 0.002$, ****, $p < 0.0001$). C) qPCR data for *GFP* mRNA levels using different concentrations of auxin (3 biological replicates). Values were normalised to housekeeping gene *RpL4* (*Ribosomal Protein L4*) and relative expression levels were calculated using the ΔΔCt method. Y-axis displaying ΔΔCt values and statistics done using Ordinary one-way ANOVA (**, $p < 0.005$). See Figure 2 - source data 1 for raw data.

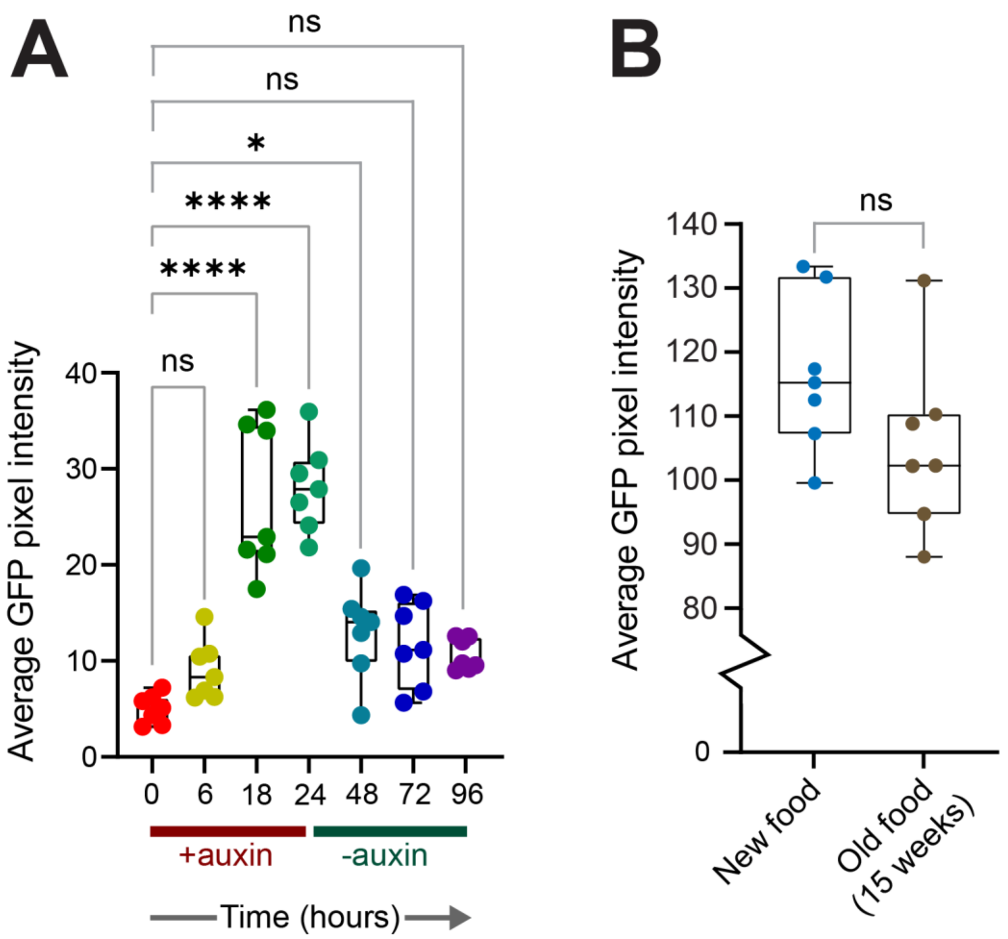


**Figure 2- figure supplement 2. On-off dynamics of AGES in adult flies and stability of auxin fly food.** A) Quantification of GFP levels in female abdomens (expressed in the fat body). Pixel intensity thresholding was performed to isolate abdomens as regions of interest. The average pixel intensity intensities were quantified and analysed using Ordinary one-way ANOVA (*, $p < 0.05$, ****, $p < 0.0001$). B) Quantification of GFP levels in female abdomens with freshly made auxin food and 15-week old food. See Figure 2 - source data 1 for raw data.

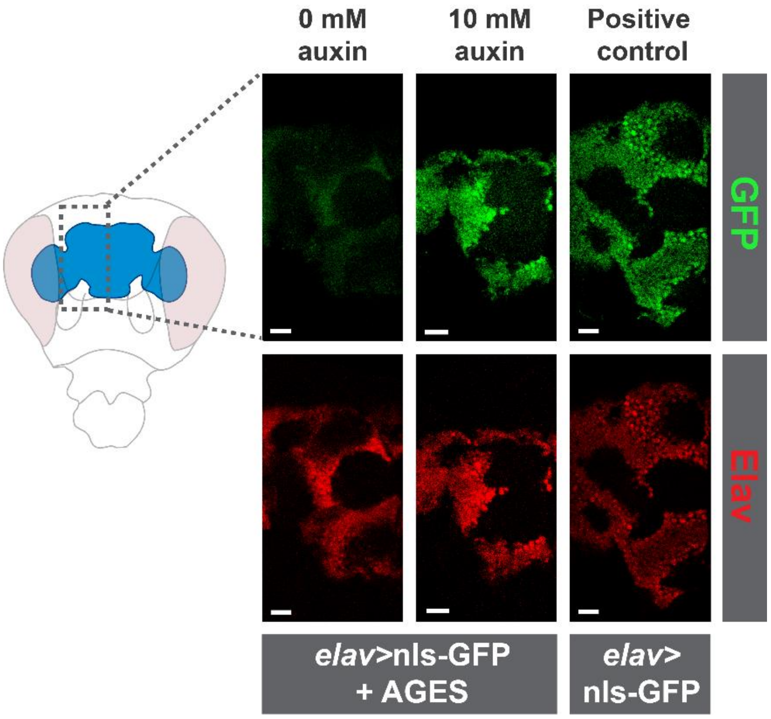


**Figure 4 – figure supplement 1. AGES allows induction of pan-neuronal GAL4 activity in the adult brain.** Confocal images of adult brains stained with anti-GFP and anti-Elav. Adult were fed food containing 5 mM auxin for 24 hours. Scale bars represent 20 µm.

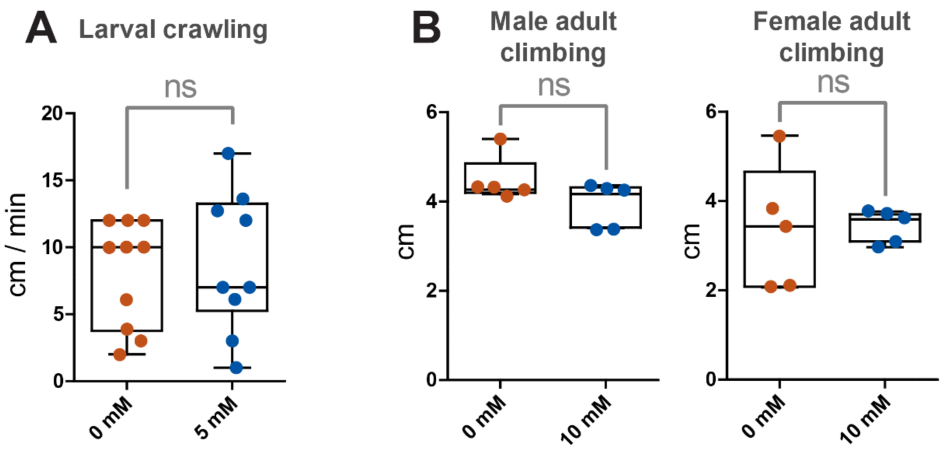


**Figure 5– figure supplement 1. Working concentrations of auxin do not impact locomotor function of wild type *Drosophila*.** A) Larval crawling speed on 0 mM food (10 larvae) and 5 mM auxin food (10 larvae). B) Distance climbed in climbing assay for males and females on 0 mM food (5 separate vials and a total of 144 flies for males and 130 flies for females) and 10 mM auxin food (5 separate vials and a total of 141 flies for males and 141 flies for females). Analysed using a paired t-test. See Figure 5 - source data 1 for raw data.

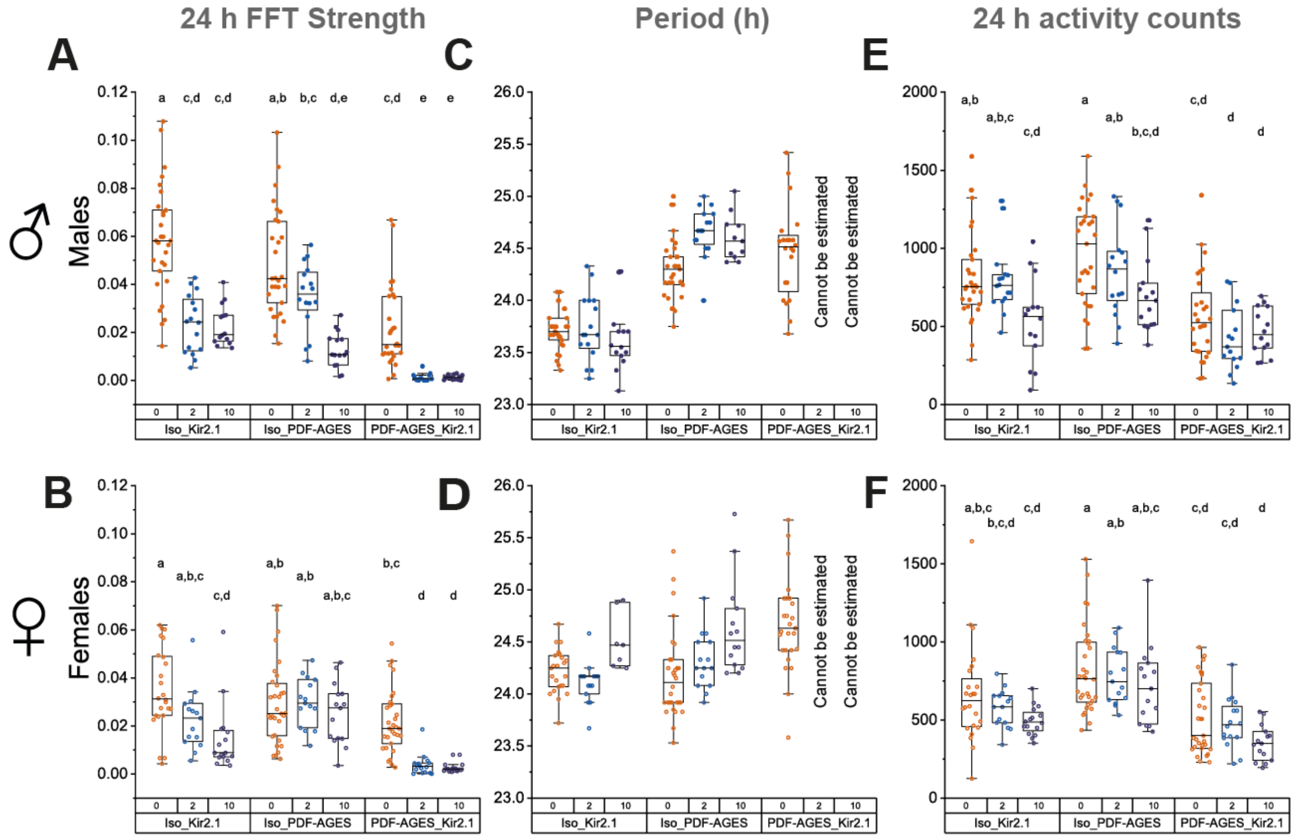


**Figure 6 – Supplement 1. Dose-dependent NAA effects on behaviour of AGES parental controls.** Behavioural data for *PDF*-GAL4; AGES> *UAS*-Kir2.1 flies and their parental controls on standard food (orange), 2 mM NAA (blue) or 10 mM NAA (purple). A, B) 24-hour FFT power on DD days 2-8 for male (A) and female (B) *PDF*-GAL4; AGES> *UAS*-Kir2.1 flies. 2 mM NAA data is replotted from Fig. 6. Means were compared by two-way ANOVA by genotype and food substrate. Means sharing the same letter are not significantly different from one another by Tukey's *post hoc* test ($p > 0.05$). C, D) Period length for male (C) and female (D) flies. Means were compared by Student's *t*-test for flies of the same genotype on different food substrates. E, F) Average 24-hour activity counts for male (E) and female (F) flies. Means were compared by two-way ANOVA by genotype and food substrate. Means sharing the same letter are not significantly different from one another by Tukey's *post hoc* test ($p > 0.05$). See Figure 6 - source data 1 for raw data and *p*-values.

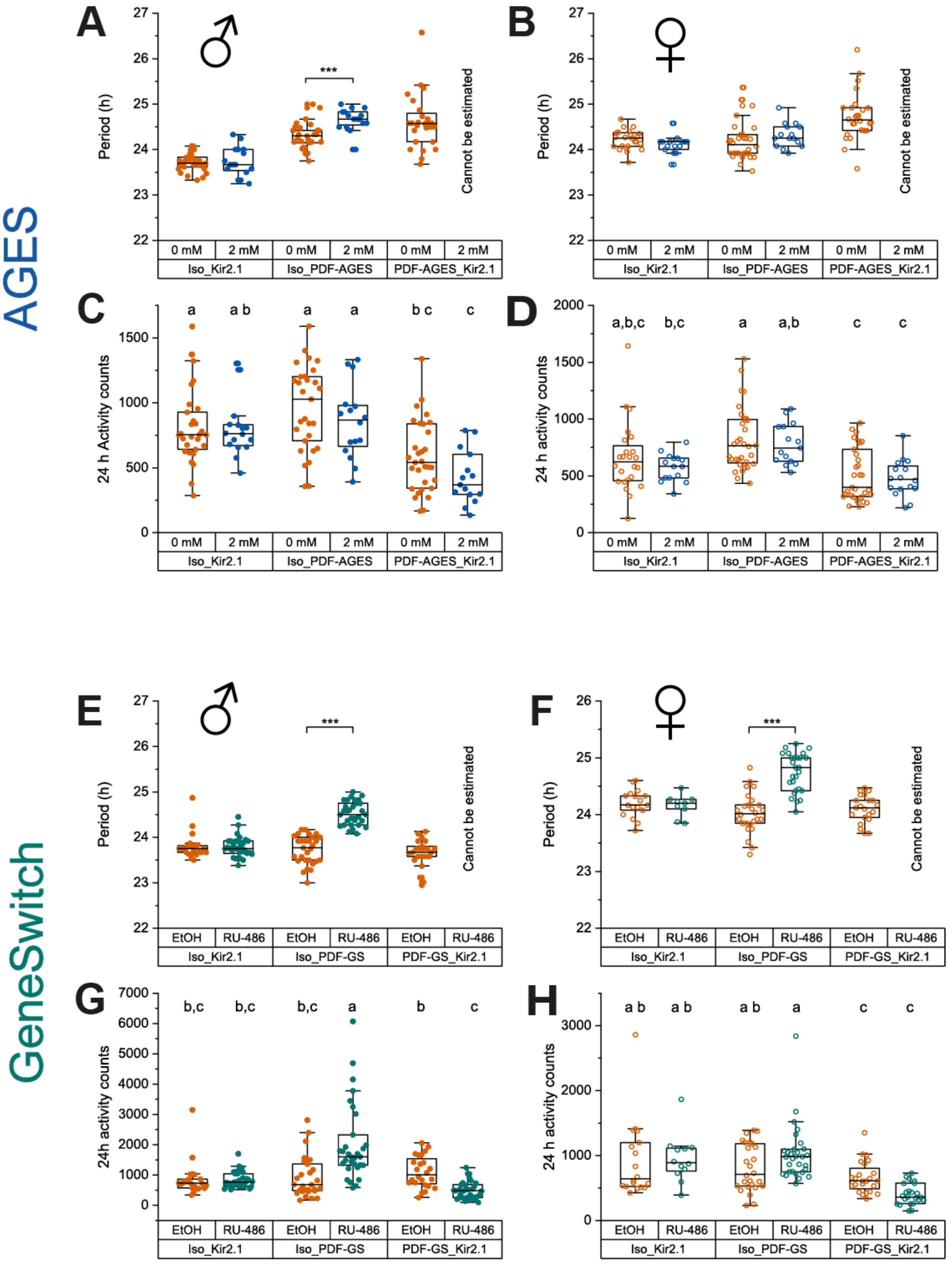


**Figure 6 – Supplement 2. Effects of AGES and GeneSwitch induced expression of Kir2.1 in PDF+ clock neurons on circadian period length and average 24h locomotor activity.** A, B) Period length estimated by chi-squared periodogram on days 2-8 of DD for male (A) and female (B) *PDF*-GAL4; AGES> *UAS*-Kir2.1 flies and their parental controls on standard food (orange) and food supplemented with 2 mM NAA (blue). Means were compared by Student's *t*-test for flies of the same genotype on different food substrates. C, D) Average 24-hour activity counts on days 2-8 of DD for male (C) and female (D) *PDF*-GAL4; AGES> *UAS*-Kir2.1 flies and their parental controls on standard food (orange) and food supplemented with 2 mM NAA (blue). Means were compared by two-way ANOVA by genotype and food substrate, and Tukey's *post hoc* test. Only genotype had significant effects on activity ($p$ = 2.86 x $10^{-7}$) and there was no significant interaction effect. Means sharing the same letter are not significantly different from one another by Tukey's *post hoc* test ($p > 0.05$). E, F) Period length for male (E) and female (F) male *PDF*-GAL4-Geneswitch>*UAS*-Kir2.1 flies and their parental controls maintained on vehicle control food (orange) and food supplemented with 466 mM RU-486 (teal). Statistics as in panels A and B. G, H) Average 24-hour activity counts for male (G) and female (H) *PDF*-GAL4-Geneswitch>*UAS*-Kir2.1 flies and their parental controls maintained on vehicle control food (orange) and food supplemented with 466 mM RU-486 (teal). Statistics as in panels C and D. Genotype had significant effects on activity in both males and females ($p$ = 4.48 x $10^{-7}$, $p$ = 3.60 x $10^{-2}$, respectively) and there was a significant interaction between the effects of genotype and food substrate in both males and females ($p$ = 0.007, $p$ = 3.76 x $10^{-9}$, respectively). See Figure 6 - source data 1 for raw data and $p$-values.

**Fly food recipe**

12 litres of Tap water

240 g Polenta

96 g Agar

1200 g Brewer's Yeast (ACROS Organics)

960 g Fructose

60 ml Nipagin (15% in ethanol)

90 ml Propionic acid

1. Add the polenta, agar, and 10 litres of tap water to the pot.

2. Bring to a boil on a high heat stirring continuously.

3. Premix the yeast and fructose to prevent clumps.

4. Once it is boiling, turn the heat down to a minimum and add the yeast and fructose.

5. Simmer for 10 minutes on low heat.

6. Turn off heat and add the remaining 2 litres of water.

7. Leave to cool, to around 70$^{o}$C before adding Nipagin and propionic acid (and auxin if required).

8. Aliquot into vials, bottles or plates.
